# Clustering the Planet: An Exascale Approach to Determining Global Climatype Zones

**DOI:** 10.1101/2023.06.27.546742

**Authors:** Jared Streich, Anna Furches, David Kainer, Benjamin J. Garcia, Piet Jones, Jonathon Romero, Michael R. Garvin, Sharlee Climer, Peter E. Thornton, Wayne Joubert, Daniel Jacobson

**Affiliations:** Oak Ridge National Laboratory, Computational and Predictive Biology, Oak Ridge, TN, USA; The Bredesen Center for Interdisciplinary Research and Graduate Education, University of Tennessee, Knoxville, TN, USA; University of Missouri-St. Louis, St. Louis, MO, USA

**Keywords:** Exascale Performance, Parallel Algorithms, GPUs, Vector Similarity Metrics, Geoclimate Zones, Climate Classification

## Abstract

We present an exascale approach for producing global scale, high resolution, longitudinally based geoclimate classifications. Using a GPU implementation of the DUO Similarity Metric on the Summit supercomputer, we calculated the pairwise environmental similarity of 156,384,190 vectors of 414,640 encoded elements derived from 71 environmental variables over a 50-year time span at 1km^2^ resolution. GPU matrix-matrix (GEMM) kernels were optimized for the GPU architecture and their outputs were managed through aggressive concurrent MPI rank CPU communication, calculations, and transfers. Using vector transformation and highly optimized operations of generalized distributed dense linear algebra, calculation of all-vector-pairs similarity resulted in 5.07 × 10^21^ element comparisons and reached a peak performance of 2.31 exaflops. We demonstrated this method using existing and synthesized climate layers to show how geography can be parsed using high-performance computing. Geoclimate zones are important tools for understanding how environmental variables impact natural systems, particularly for agriculture and conservation with relevance to climate change. Historically, classification systems have been low resolution, based on limited variables, or subjective. To identify climate classes, we clustered DUO outputs at varying stringency, producing 69, 133, 340, and 717 global geoclimate zones. Our approach produced global scale, high resolution, longitudinally informed climate classifications that can be used in precision agriculture, cultivar breeding efforts, and conservation programs.

## 1. Introduction

Climate patterns vary across space and time and are created by numerous environmental factors that impact natural systems. As current agricultural lands become less productive due to changing climate and pathogen distributions [1, 2], there is an increasing need for modernized, fine-scale, high-resolution climate zone delineations that allow precision matching of extant plants to appropriate environments and inform cultivar engineering efforts for marginal lands.

Most common classification methods are based on the Köppen-Geiger system [3], which identifies five major climate types based on temperature and precipitation. Trewartha and Horn (1980) expanded this system to include seven major and 14 minor climate types using finer gradations of precipitation and temperature [4]. More recent efforts have typically been low resolution or relied on historical vegetation and/or zoological mapping information, often based on individual expertise and visual assessment of previous maps. Two such examples are Olson’s manual delineation of eight biogeographic realms and 14 biomes [5], and the use of 7842 species’ ranges by Echeverría-Londoño *et al*. (2018) [6] to classify North and South America at 100km^2^ resolution. In 2011, Kumar *et al*. (2011) [7] used supercomputers to perform a k-means clustering analysis on North America. However, analyses incorporated a limited number of environmental variables and did not incorporate longitudinal (across years) or temporal (within year) data.

Although these definitions addressed deficiencies of the Köppen-Geiger system, they were low resolution, calculated from a limited number of variables, or based on intrinsically biased human-designated thresholds. By using high performance computing to analyze complex, longitudinal global scale geoclimate data sets, the accuracy and resolution of currently defined geoclimate zones can be greatly increased. Doing so makes it possible to better match crops with suitable local agricultural conditions or identify target traits for breeding and engineering programs.

Here, we capture daily climate patterns using 71 environmental variables over a 50-year time span at 1km^2^ resolution across the planet. After uniquely encoding data into 156,384,190 climatype vectors containing 414,640 elements each, we used dense linear algebra operations to calculate correlation values by performing an all-versus-all pairwise comparison. To do so, we implemented the DUO Similarity Metric [8] on GPU Tensor Cores in the Summit supercomputer at the Oak Ridge National Laboratory to perform 5.07 × 10^21^ element comparisons, achieving a peak performance of 2.31 exaflops. The resulting correlation values were modeled as a network and clustered at various granularities to identify high-resolution climate zones across the globe.

Our method facilitates rapid, accurate, unbiased identification of climate zones and environmentally correlated regions. When combined with efforts to optimize plant phenotypes and environmental adaptation, this approach contributes to the advancement of precision agriculture, which will help meet growing food and energy insecurity of an expanding human population.

## 2. Materials and Methods

### 2.1. The DUO Similarity Metric

The DUO Similarity Metric algorithm calculates the similarity between two vectors by binning matrix values into low or high (0 or 1) categories and looking for regions of the vectors that correlate [9, 8, 10]. Comparisons between vector *A* and vector *B* produce four possible combinations: A_high_ vs B_high_ (or “HH”); A_high_ vs B_low_ (“HL”); A_low_ vs B_high_ (“LH”); and A_low_ vs B_low_ (“LL”). The algorithm is represented by the equation:

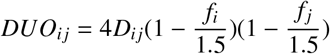

For every *ij* pair where *i* ϵ {*A*_*H*_, *A*_*L*_} and *j* ϵ {*B*_*H*_, *B*_*L*_} and where *D*_*ij*_ is equal to the fraction of vector lengths of when *A* and *B* co-occur. The values *f*_*i*_ and *f*_*j*_ are the fraction of *i* and *j* vector cells in *A* and *B* respectively. This algorithm calculates the correlation (and effectively, anti-correlation) values between all compared vectors and accounts for frequency effect by scaling the resulting values according to the fraction of high/low values in vectors being compared.

### 2.2. Data Extraction

Geoclimate data collected from 156,384,190 square kilometers of dry land (excluding Antarctica) were obtained from www.worldclim.org [11], www.earthenv.org [12, 13], www.fao.org [14], www.cgiar-csi.org [15, 16], and the Global High-Resolution Soil-Water Balance data set [17] (Figure 1). Data extraction from GeoTiffs was performed using the Geospatial Data Abstraction Library (GDAL) [18]. For cases in which GeoTiffs were not available, in-house code was used to generate data layers. We calculated unique dissipation rates as linear models for each of 48 spectral bands of light. Other layers produced by in-house code included soil nutrient layers for which we calculated ion activity based on temperature, soil pH, and soil water dilution for the top 12 most important nutrients (nitrogen, boron, calcium, phosphorus, sulfur, potassium, zinc, copper, molybdenum, manganese, magnesium, and iron).

**Figure 1:**
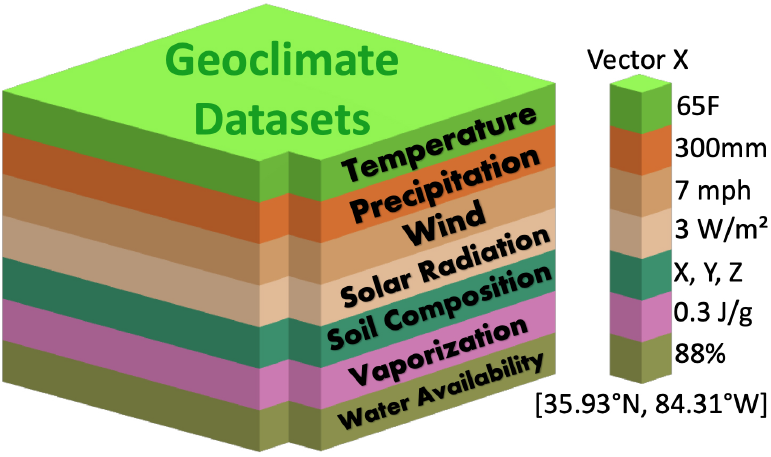
Example schematic of a subset of geoclimate datasets used to extract vectors for each square kilometer of dry land on the planet. Each set of geoclimate data contains averaged information for a specific month or year.

From each of the final 71 geoclimate layers, fifty years’ worth of historical monthly-averaged data was extracted and interpolated into 365 days’ worth of data per layer, resulting in ≈3,618,391,000,000 observation points.

### 2.3. Vector Encoding

DUO bins values into 1s and 0s; however, climate variable distributions are rarely binomial and vary considerably from equatorial to polar locations. To convert complex variables into binary data, we used a bit-based encoding method and expanded a 0/1 input into to a user-defined range of bits (Figure 2). For each location and geoclimate variable, we converted the daily numeric geoclimate values to binary, then concatenated daily data into a single year-long vector. Finally, the year-long vectors for each climate variable were concatenated to produce a full length binary encoded vector.

**Figure 2:**
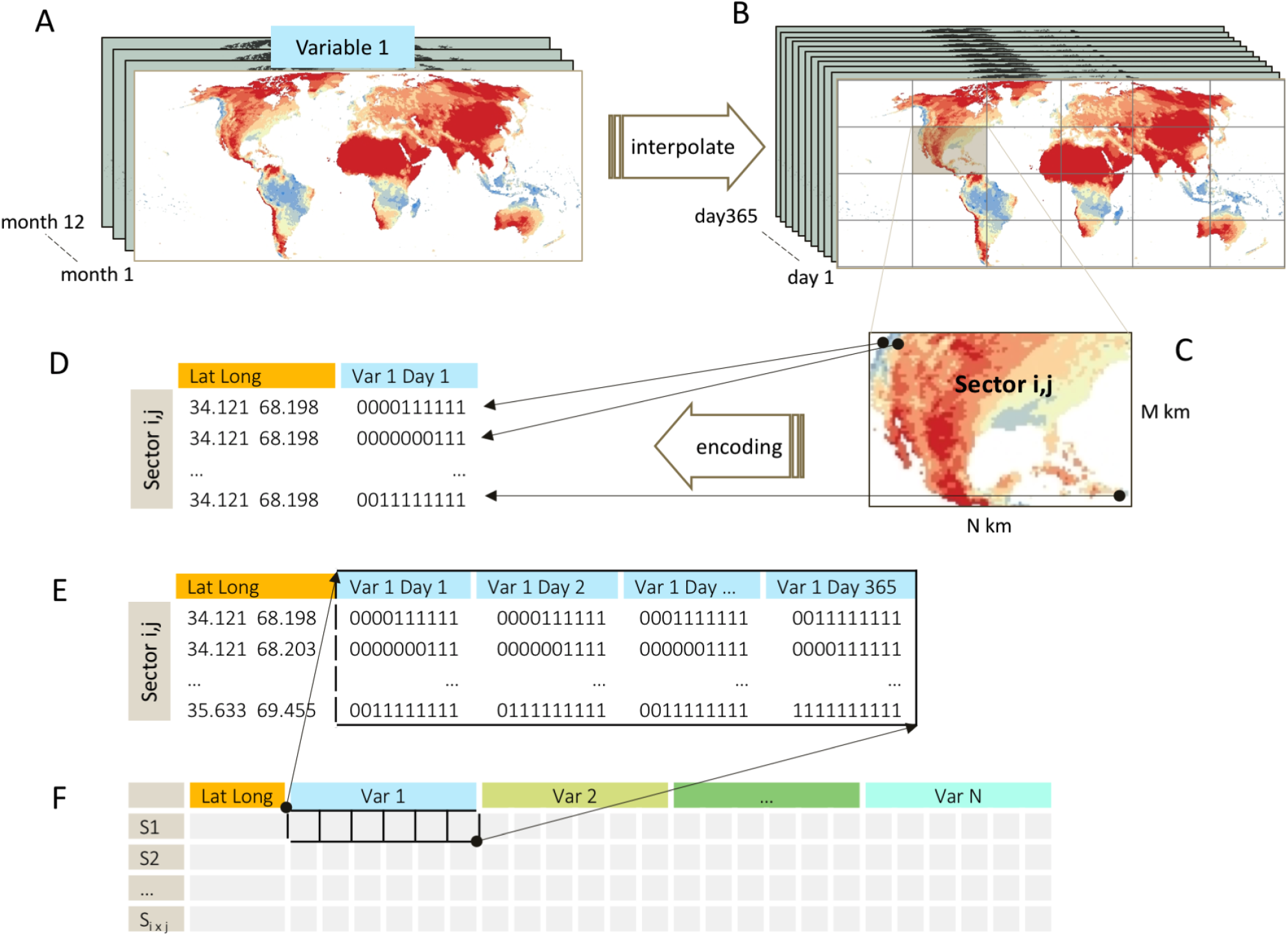
Multiple global geoclimate data sets (variables) were obtained and encoded in 150 million binary vectors for a pairwise correlation analysis using DUO. A) Each variable initially consisted of TIFF image data, with 12 images representing monthly averages over 10, 30, or 50 years depending on the data set. B) Monthly data was interpolated to produce 365 days’ worth of data per variable. C) For parallelization, the world was divided into sectors. Within a sector(*i,j*), for each day of the year the numeric variable value at every latitude and longitude was extracted and converted into a binary vector using bit-based binary encoding. (D) Full length vectors were formed by concatenating the daily vector data for all variables (E, F), thus representing all available climate/environmental data at every latitude/longitude point on Earth as a vector.

### 2.4. Large Data Set Clustering

DUO output was thresholded at 0.80 to identify highly similar geographical locations. DUO produces four similarity values per pair of vectors (HH, HL, LH and LL); positive associations (HH or LL >0.80) were sub-sampled and clustered using MCL with inflation values of 1.4, 2.0, 3.0, and 4.0 to find clusters of environmentally similar land at different granularities [19].

### 2.5. Compute System

Summit consists of 4,608 compute nodes, each with two 22-core IBM POWER9 processors and six Nvidia Volta V100 GPUs connected to a POWER9 by NVLINK-2 with 100 GB/sec bidirectional peak performance. V100 GPU peak double precision performance is ≈7 TF, theoretical peak mixed precision performance is ≈125 TOps, and peak memory bandwidth is 900 GB/sec. Each node contains 512 GB main memory and each GPU contains 16 GB HBM2 memory. The nodes are connected by Mellanox Infiniband fat tree interconnect with adaptive routing and equipped with a 1.6 TB NVMe burst buffer device.

Summit is connected to the GPFS file system Alpine; software versions were GCC 6.4.0, MAGMA 1.6.2, Spectrum MPI 10.2.0.11-20190201 and CUDA 9.2.148. The `jsrun` tool was used for application launch, with six MPI ranks per node each with 1 GPU and 7 OpenMP threads mapped to CPU cores. Environment variables `PAMI_IBV ENABLE DCT=1`, `PAMI_ENABLE_STRIPING=1`, `PAMI_IBV_ADAPTER _AFFINITY=0`, `PAMI_IBV_QP_SERVICE_LEVEL=8`, `PAMI_IBV_ENABLE_OOO_AR=1` were set to use adaptive routing and to limit per-node storage needed for MPI buffers.

### 2.6. Vector Handling, GEMM and MAGMA

Calculating similarity of *x* vectors of length *y* requires *O*(*x*^2^*y*) of *O*(*xy*) inputs. We used GEMM BLAS-3 dense linear algebra matrix-matrix product operations to directly exploit structural similarity calculations of vectors [20]. On top of GEMM based operations in a hybrid architecture, we adapted the MAGMA library to speed similarity calculations [21]. DUO also required modification for operations at bit-level: mixed precision modifications for GEMM and Tensor Core methods; we refer to this tensor core implementation as DUO-TC.

### 2.7. Hardware Optimization

We used CUDA intrinsics for fast hardware instructions and the `CUDA _ _popcll` intrinsic to count the 1 bits within a word [22, 20]. We used the `cublasGemmEx` function on Summit’s NVIDIA V100 GPU to increase performance of the DUO algorithm through use of Tensor Core hardware and mapping DUO input vectors to half precision FP16 numbers then applying a matrix-matrix product.

### 2.8. Load Balancing and Redundant Computations

To reduce redundant computations from data symmetries while maintaining load balance, subsets of computed values were chosen carefully. Three main tasks were used to reduce redundancy: unique metrics are computed at least once, but with as few repetitions as possible; proper load balancing across MPI ranks; within rank data is chosen where the modified GEMM operations are as large as possible and, ideally, of the same length as the vectors (for high-performance computing context) [23].

### 2.9. Parallelization

Efficiently distributed dense linear algebra usually requires multidimensional parallelism [24]. Here we use three parallelism axes: set partitions of vectors, decompose said vectors by length, and replicate vectors to distribute computation of result blocks. Each of the 4608 nodes on Summit has 512 GB of memory per node. Because the input file containing 156,384,190 vectors of 414,640 elements was 118 TB (7.4 TB when binarized), input data had to be distributed across all Summit nodes. This large pairwise comparison required enormous intranodal communication; rapid communication between nodes required a novel stepwise scheduling in which partitions of vectors were compared by those used by another offset MPI rank.

### 2.10. Performance Measurement

Our measurement methodology was described in detail in [25]. In brief, timings were collected using the `gettimeofday` function, using `cudaDeviceSynchronize` and MPI barriers to synchronize the nodes. The GEMM operation counts were computed analytically within the code. Unless otherwise specified, the core algorithm computation time was measured without I/O. Time for input from the Alpine file system was typically small as a fraction of runtime. Time for output to the burst buffers was highly dependent on the output threshold factor used for the run, but this setting was calibrated to minimize the impact of output time cost to the code.

## 3. Results and Discussion

### 3.1. GPU Kernel Results

Each node of Summit contains 6 NVIDIA V100 GPUs which can execute FP16, up to 86 teraFLOP/s through PCIe. Due to the strategic exploitation of the V100 Tensor Cores, a scalable performance of 2.15 exaflops was achieved for the 4550 nodes included in a scaling study in (Figure 3). In a separate optimized run, a peak performance of 2.31 exaflops was accomplished.

**Figure 3:**
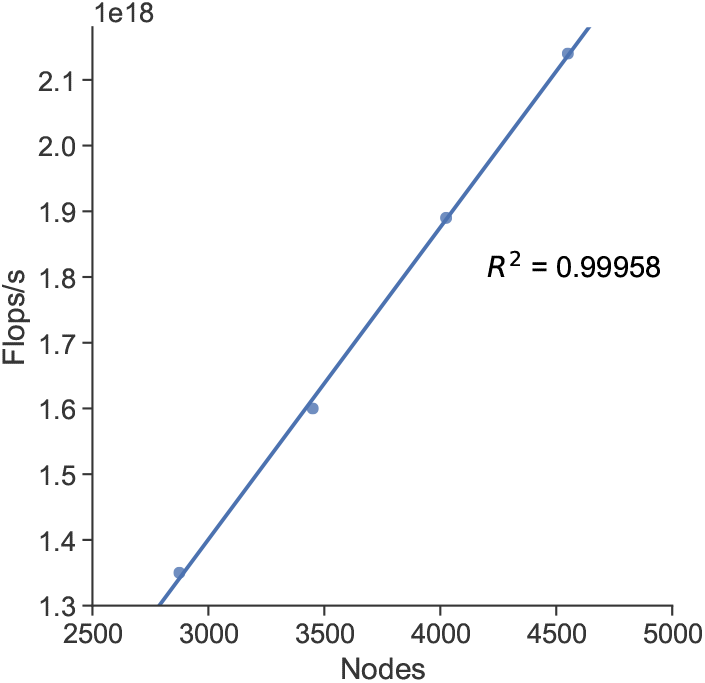
Parallel execution flops rate.

### 3.2. Runtime Components

The core computation time in Table 1 represents the time for calculating DUO metrics and the time for initializing the vectors and computation of metrics for one phase. Data input takes considerable time due to the size of the input.

**Table 1:** Timings in seconds of execution components for different numbers of nodes.

| Number of Nodes | 2875 | 3450 | 4025 | 4550 |
| --- | --- | --- | --- | --- |
| Core Computation | 30.41 | 25.08 | 21.80 | 19.15 |
| Vectors Initialization | 0.72 | 0.66 | 0.59 | 0.50 |
| Metrics Initialization | 1.44 | 1.14 | 0.96 | 0.84 |
| Input | 26.87 | 34.31 | 42.21 | 50.33 |

### 3.3. Strong Scaling Results

We used the strong scaling method to scale to a larger number of processors. We fixed the problem size to 156M vectors of 414K elements and increased the number of computing nodes from 2875 to 4025 nodes. This range was chosen and set to use adaptive routing due to the large input; a lower memory limit of 2300 nodes and limited per-node storage needed for MPI buffers could not accommodate the input on the GPUs, where the main computations were performed. The parallel execution flop rates for strong scaling tests ranged from 1.35 to 2.13 exaflops/s (Figure 4) as the number of nodes increased. The clear, linear, strong scaling bodes well for using the DUO-TC solution at scales of data even larger than the one in this study.

**Figure 4:**
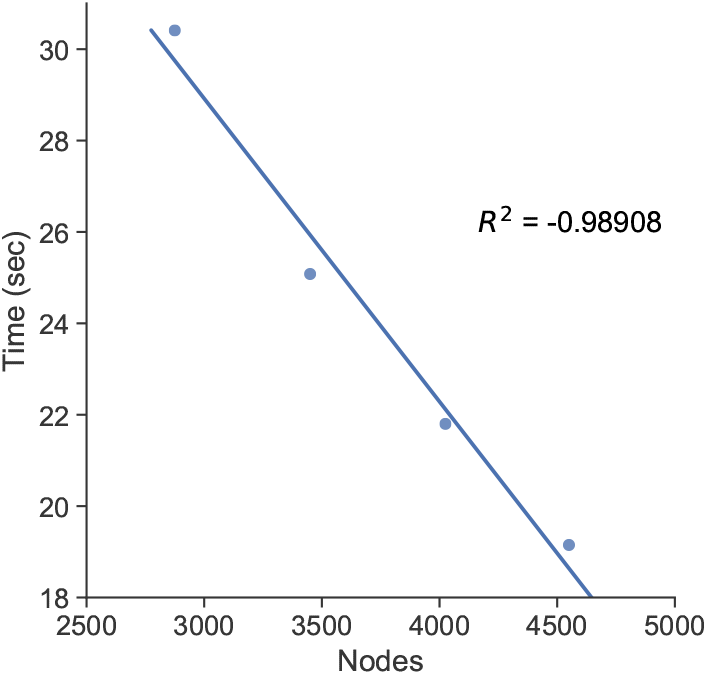
DUO-TC strong scaling.

### 3.4. Weak Scaling Results

We performed a weak-scaling benchmark of DUO on Summit, maintaining a fixed problem size per GPU, and calculating the number of comparisons per GPU per second to observe how the number of comparisons per GPU per second varied over the number of nodes (Figure 5). Improved scalability was observed when using 3450 nodes or more, with the performance further improved up to 9.77E+12 comparisons per GPU per second for 4550 nodes.

**Figure 5:**
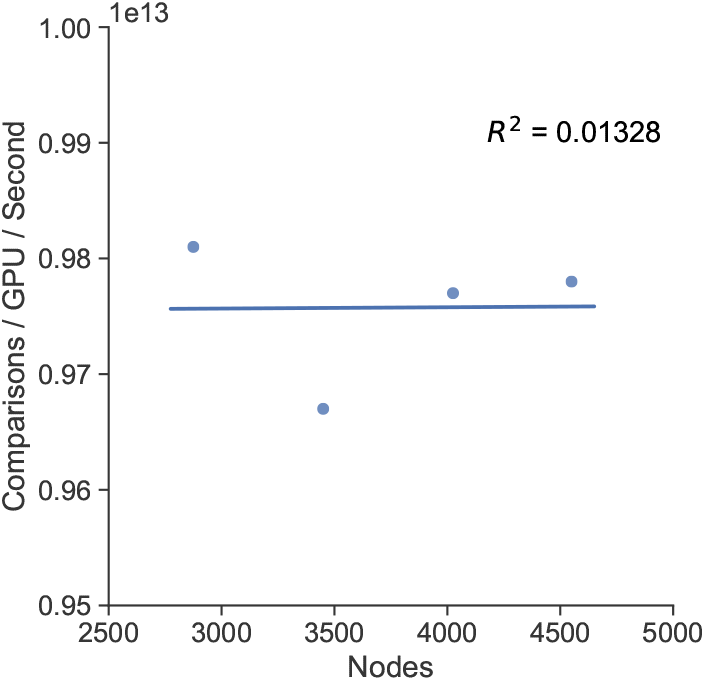
Weak scaling comparison rates.

### 3.5. Geoclimate Clustering and Correlation Results

Markov clustering of the thresholded DUO-TC outputs assigned every 1km^2^ of land on Earth to an unsupervised geo-climate zone (cluster). Note that the clusters in Figure 6 are limited to 57 colors because the human eye is unable to visualize the full data (this is roughly 1/14 of the true number of colors that would be required to represent all climate regions). Additionally, only a sub-sample of the thresholded DUO results were used as input to MCL.

**Figure 6:**
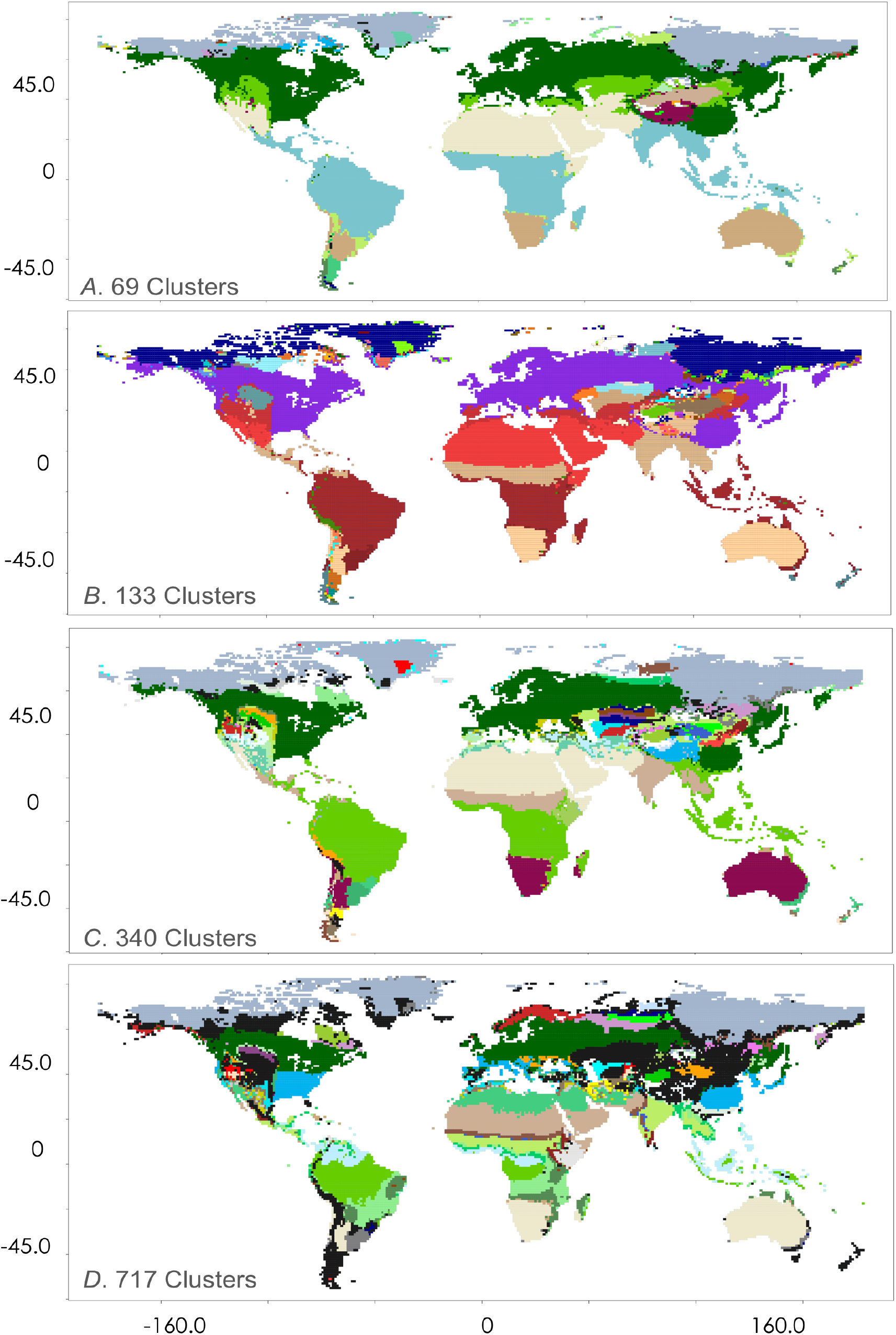
Clustering of 28,450 elements with four different inflation values. Progressively increasing MCL inflation values increases granularity of the geographic distribution of clusters. Many clusters are common across geography, while others are rare and are colored dark grey

The gradients of global climate zones become clearer and more refined by examining larger inflation values (and therefore higher granularity of clusters). Increasing to 717 clusters identifies many correlated regions from distant positions on the planet at an increasingly fine scale.

## 4. Conclusions

We performed global, high resolution climate classification using a novel implementation of DUO capable of performing calculations at an improved rate for very large data sets. We calculated pairwise similarity at exascale using a unique vector encoding method to capture complex, longitudinal geoclimate data. We provide a comprehensive, mathematical representation of the Earth’s correlated environments using network theory. No other system currently exists that approaches the resolution of climate clustering we present here. In combination with explainable-AI-based genomic selection and bio-engineering methods, this would provide the ability to optimize cultivars adapted to specific environments and at a global-scale.

The results presented here demonstrate the combined power of DUO-TC, MCL, and supercomputing to partition large data into refined groups even for problems on a global scale. Furthermore, the methods we present are highly adaptable; many data types can be encoded and analyzed using our DUO-based methods. The same approach can be applied to demographic, geological, agricultural, and ecological data sets. DUO outputs can also be analyzed with several different clustering algorithms such as Breadth First Search (BFS), k-means, or MCL.

Continued work in this field will include the addition of more layers, such as pollution, population density, CO_2_ levels, and green areas in urban environments. A wealth of data will soon become available from technological advances in remote sensing (*e.g*. drones, low power monitoring devices, new satellite platforms). Oceanographic information such as bathymetry, benthic habitat, sea surface temperature, and distribution of large oceanic currents can also be added to oceanic regions as well. The integration of more refined temporal components and dynamic environmental and species composition across years will also improve the accuracy and utility of climate zones.

## Credit authorship contribution statement

**Jared Streich:** Conceptualization, Methodology, Software, Formal Analysis, Investigation, Data Curation, Writing - Original Draft, Writing – Review & Editing, Visualization. **Anna Furches:** Formal Analysis, Writing - Original Draft, Writing – Review & Editing, Visualization, Project Administration. **David Kainer:** Methodology, Software, Writing – Original Draft, Writing – Review & Editing, Visualization. **Benjamin J. Garcia:** Formal Analysis, Writing - Original Draft. **Piet Jones:** Formal Analysis. **Jonathon Romero:** Formal Analysis. **Michael R. Garvin:** Writing - Original Draft. **Sharlee Climer:** Software, Writing - Review & Editing. **Peter E. Thornton:** Methodology, Writing - Review & Editing. **Wayne Joubert:** Software, Writing - Review & Editing. **Daniel Jacobson:** Conceptualization, Resources, Writing - Original Draft, Writing - Review & Editing, Supervision, Funding Acquisition.

## Declaration of Competing Interest

The authors declare no competing interests.

## Acknowledgements

The authors would like to thank João Gabriel Felipe Machado Gazolla - without whom this work would not have been possible, Jack Wells, and Don Maxwell for their assistance; we thank Ramanuja Simha, Mikaela Cashman, John Lagergren, and Matthew Lane for valuable discussions. This research used resources of the Oak Ridge Leadership Computing Facility Summit Early Science Program and was supported by the Center for Bioenergy Innovation, the Plant-Microbe Interface SFA, the Feedstock Genomics Program (all supported by the Office of Biological and Environmental Research in the DOE Office of Science) and LDRD #8321 at the Oak Ridge National Laboratory, which is managed by UT-Battelle, LLC, for the US Department of Energy under contract DE-AC05-00OR22725.

## Notes

### Competing Interest Statement

The authors have declared no competing interest.

